# A qualitative mathematical model of immunocompetence with applications to SARS-CoV-2 immunity

**DOI:** 10.1101/2021.12.08.471857

**Authors:** Javier Burgos-Salcedo

## Abstract

A qualitative mathematical model of the notion of immunocompetence is developed, based on the formalism of Memory Evolutive Systems (MES), from which, immunocompetence is defined as an emergent structure of a higher order arising from the signal networks that are established between effector cells and molecules of the immune response in the presence of a given antigen. In addition, a possible mechanism of functorial nature is proposed, which may explain how immunocompetence is achieved in an organism endowed with innate and adaptive components of its immune system. Finally, a practical method to measure the immunocompetence status is established, using elements of the theory of small random graphs and taking into account the characteristics of the immune networks, established through transcriptional studies, of patients with severe COVID-19 and healthy patients, assuming that both types of patients were vaccinated with an effective biological against SARS-CoV-2.

## Introduction

Defending oneself against pathogens is not something that should be taken lightly, especially now, when the COVID-19 pandemic affects practically human populations worldwide, and the quest for those mechanisms involved in the immune response against the SARS-CoV-2 virus deserves a lot of attention (Chowdhury 2020; Mortaz 2020; Alturaiki 2021). Traditionally, the concept of immunocompetence, understood as a measure of the ability of an organism to minimize the fitness cost of infection via any means after controlling for previous exposure to appropriate antigens (Owens and Wilson, 1999), has been the object of research by evolutionary biologists, and has been widely used for the genetic improvement of species of economic importance, such as poultry, and also, for the study of the natural immunity in wild animals (Viney 2005; Ask 2007).

The mammalian immune system is a dynamic multiscale system composed of a hierarchically organized set of molecular, cellular, and organismal networks that act in concert to promote effective host protection against pathogens. These networks range from those involving gene regulatory and protein-protein interactions underlying intracellular signalling pathways, and single-cell responses, to increasingly complex networks of in vivo cellular interaction, positioning, and migration that determine the overall immune response of an organism (Subramanian 2015). Immunity is thus not the product of simple signalling events but rather nonlinear behaviours arising from dynamic, feedback-regulated interactions among many components.

Taken into account the structure of the immune system, in this paper a model for immunocompetence is proposed using the mathematical formalism of Memory Evolutive Systems (MES), first introduced by Ehresmann and Vanbremeersch (2007) to develop mathematical models for evolutionary, multi-scale, multi-temporality and self-organized systems. In the present paper, we construct a notion of immunocompetence as a hyperstructure emerging from the cooperation of the components of a pattern, in this case, the effector immune cells, which are devoted, in principle, to carry out a particular task e.g., to recognize and neutralize a particular antigen.

Once the notion of immunocompetence has been defined, we use it to propose a possible mechanism of action by which effective vaccines against COVID-19 can act as transition drivers between the immune configurations of an individual towards the construction of a protective immune response against the coronavirus. Finally, and as a result of defining the underlying immune pattern as a small random graph, it is possible to estimate the probability that an individual is immunocompetent to a given antigen, using information from the immune cell networks involved in the response to SARS-CoV-2 in patients with severe COVID-19 and healthy patients.

### Theoretical framework

#### Memory Evolutive Systems and Categories

Memory Evolutive Systems (MES), developed by Ehresmann and Vanbremeersch (2007), represents a mathematical model for natural open self-organizing systems, such as biological, sociological, or neural systems. In these systems, the dynamics are mixed with the global hierarchical memory. The MES proposes a mathematical model for autonomous evolutionary systems. It provides a framework to study and possibly simulate the structure of “living systems” and their dynamic behavior. MES is based on a conceptual mathematical domain named Category Theory (Lawvere 1997, Beurier 2019), providing a relational approach for studying evolutionary, multi-scale, multi-temporality, and self-organized systems (Ehresmann and Vanbremeersch 2019).

A category is made up of a class of objects together with a class of maps –a type of processes, or paths– on those objects, conforming a graph, here named the underlying graph, and a composition law defined over the graph following associativity and identity axioms. The category concept is intended to capture - emphasizing the concept of relationship (of application), rather than of element and belonging - the essence of a class of objects, which are related through applications, the maps in the category in question. At an increasing level of complexity, it is possible to think about how two categories can interact with each other. In mathematics, the term functor is used in the theory of categories to designate that function from one category to another that takes objects to objects and maps to maps, so that the composition of maps and identities are preserved. A functor is a term used by formal logic to designate any logical or connective operator (such as conjunction, disjunction, etc.) so that, having no semantic content, they are used to establish relationships between content.

Finally, in category theory, adjointment is a relationship that can have two functors. Two functors found in this relationship are known as adjoint functors, one is the left adjoint and the other is the right adjoint. In mathematics, specifically in category theory, adjointment is a relationship between two functors that appears frequently throughout the different branches of mathematics and that captures an intuitive notion of solution to an optimization problem.

#### Random graphs of small order

The immune system is particularly well suited for the application of network reconstruction methods and before developing the new notion of immunocompetence (Subramanian 2015). Briefly, a graph consists of a set of elements together with a binary relation defined on the set. Graphs can be represented by diagrams in which the elements are shown as points or vertices A, B, C, etc., and the binary relation as lines, links, or arrows, joining pairs of points. These links can be of different natures: more or less invariant structural links, such as spatial relations; causal relations; energetic or informational relations; or those imposing constraints of any kind. The links can also represent labile connections, corresponding to a temporary interaction (chemical reaction). In any case, the links correspond to interactions between objects, which can be more or less extended in time. However, the true importance of graphs is that, as basic mathematical structures, they arise in diverse contexts, both theoretical and applied.

Within graph theory, a very fruitful line of research has been developed since the late 1950s, which is that of random graphs, currently described as those generated by a probability distribution, or by a random process that generates them. Random graphs were originally proposed by Erdos and Renyi in 1959, who were inspired by the growth patterns of living organisms. There is an impressive theoretical corpus on random graphs in which the main results are related to structures of infinite size, these are also valid when considering graphs of finite size, that is, the number of nodes is finite (Bollobás 2001). This has made it possible to form an object of study known as small-order random graphs (Bollobás 1985), which undoubtedly can adequately represent the dynamics of biological interaction networks, and in particular, those of the immune system since these are generally finite. In the present work, the underlying graph considered is a small small-order graph made up of the effector cells of the immune response as their objects or vertices and the signals or links, between them as their maps.

### Modeling immunocompetence

Let us consider a category K modeling an organism endowed with an effector system made up of a complex network of cells and signals that enables both, innate and adaptive immune responses, to the presence of a given antigen. This complex network here will be modeled as a small random graph, whose vertices are cells and their links correspond to transmitting messages/information in the form of effector signaling molecules between them. Moreover, if X, Y and Z represent immune effector cell types:

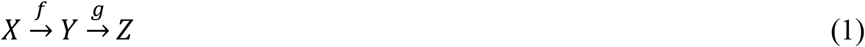

These links follow a composition law which associates to each path *(f, g)* the composite, denoted by *h: =gof*, of the path.

Inside *K*, let us consider a set or Pattern (P) of immune cells with their links or effector molecules like complement, interferons or antibodies among others (**Figure 1**). For an immune pattern (*P*), a perspective of an antigen (*A*) is a family of links (*g*_*i*_) from the epitopes of A to some components P_i_ of P (called aspects of the antigen by the immune pattern) such that any two aspects are connected inside P by a set of distinguished links (*d*). It is important to note that, to be effective, the various cells types or components of the immune pattern P, must act in a coordinated way to fight against an external attack, by the contrary, the defense strategy may fail. This coordinated action of the components P_i_ conforms a new complex, or higher, object cP in K through the action of a family *C:(c_i_)* of links such that, associated to each index I of P is a link c_i_ form the component P_i_ to cP. These family C of links are named Collective Binding Links from P to cP.

**Figure 1.**
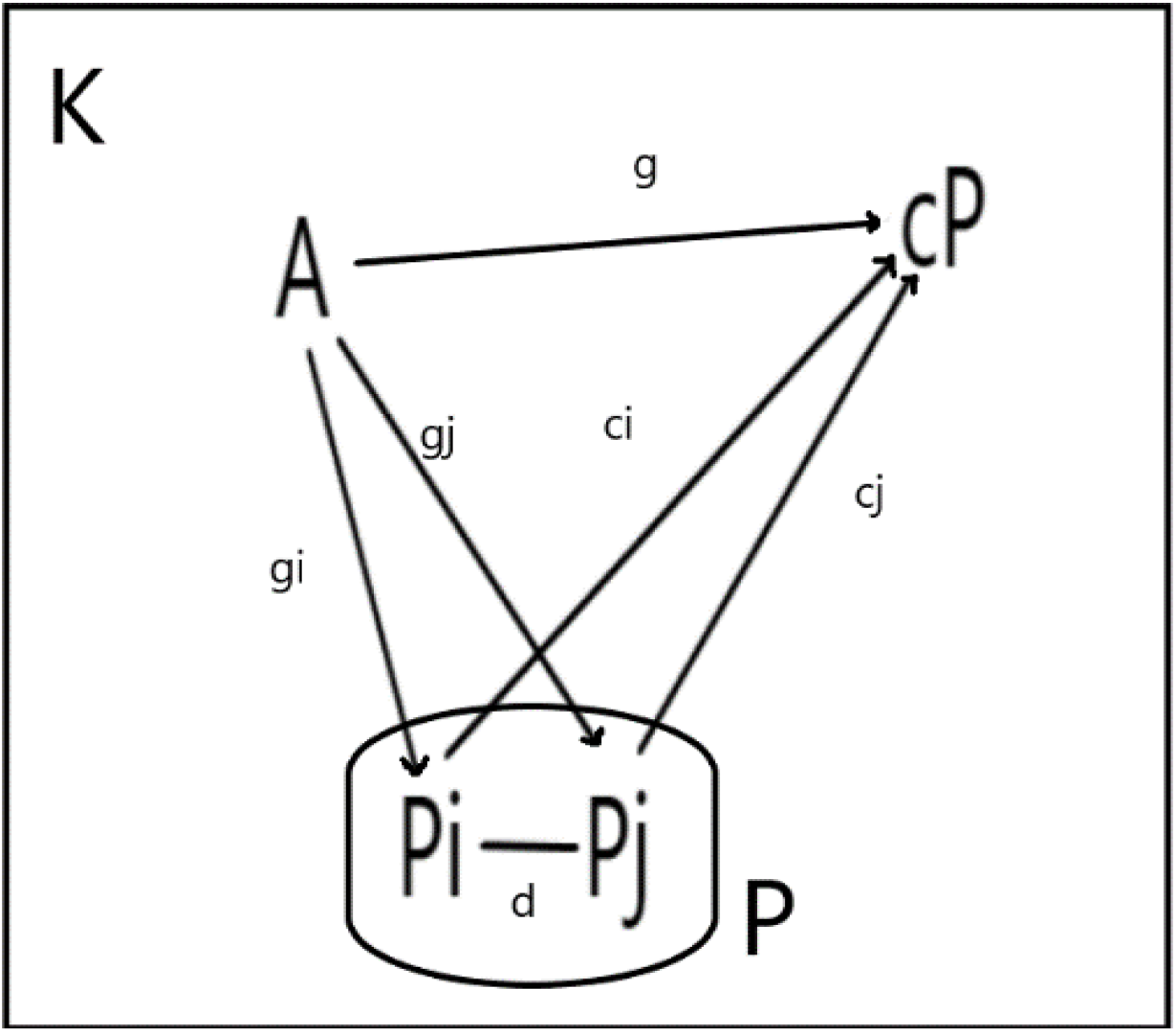
Representation of immunocompetence as an emergent hyperstructure. **Legend figure 1**. *K* represents the category of organisms with a complex immune system composed by innate and adaptative compartments. *P* represents a set of effector immune cells connected between them by signal links (*d*) as a result of antigenic stimulation (*g*_*i*_) driven by *A*. From the interaction of the subjacent network conformed by the interaction between effector cells, a set of collective binding signals or links (*c*_*i*_) emerge, generating the immunocompetent state *cP*, which can be understood as a regulatory network capable to control the proliferation of the pathogen *A*.

On the other hand, the distinguished links *d* of the immune pattern P restrains the freedom of the individual links c_i_ of the collective binding links from P to cP, which means that the larger the number of distinguished links d of the immune pattern associated with the immune response against a pathogen, the more constrains are imposed on the collective binding links, and, the smaller the number of d_i_, less constrains are imposed on the family C of collective binding links.

To introduce our notion of immunocompetence, we will need the following definition.

#### 1 Definition 1 (colimit)

An object of the category *K* is called the colimit of the pattern *P* in *K*, denoted *cP*, if the following two conditions are satisfied:

1. There exists a set of collective links (*c*_*i*_) from *P*, with components *Pi* to *Pj* connected by links *d*, to *cP*, called the collective binding links.
2. For a given pattern *P*, if the colimit exists, it is unique. This condition is usually named the universal property of colimits. The biological meaning of Condition 2 is that once established, immunocompetence acts as a central set of signaling pathways whose characteristics depends upon the perspective of the antigen A developed by those interactions between the components of P, induced by the signals g_i_ and g_j_ emitted by A. The universal property is represented by the signal *g* from *A* to *cP* (**Figure 1**).

It is important to note that a colimit corresponds with the notion of hyperstructure, and, understanding immunocompetence as the ability of the organism (*K*) to produce an optimal-protective-immune response following the exposure to an antigen *A* as a result of the synergistic actions of the components of the immune pattern *P*, we can propose the following notion:

### Immunocompetence

An organism is immunocompetent if there exists a set of collective binding links, generated between the immune effector components of the immune system as a result of the interaction with an antigen, which induce a hyperstructure (cP), conformed by a set of central signaling pathways who regulates the immune function of the organism, leading to the recognition, elimination, and memorization of the antigenic object. In other words, the coherent functional role of P induces a higher structure, conformed by a set of immunoregulatory signaling pathways, in the organism K, which is precise, the immunocompetent status of the organism.

To give an intuitive idea of the notion of immunocompetence we proposed in the present work, let us consider a useful illustration by Ehresmann and Vanbremeersch (2007): “The cooperation of the components of a pattern (P) can be temporary, as in a group of people who decide to meet to carry out a particular task. However, if it lasts for a long period, their cooperation may be reinforced, with some specialization of the roles of the different members. Finally, the group as such can take its own identity and become legalized as a professional association, binding its members and requiring that they operate coherently to fulfill the functions devoted to the association, this will lead to a hyper-structure, categorically known as the optimal object or colimit of the pattern”.

Having defined a formal notion of immunocompetence as a colimit, we are prepared to put forward a hypothesis about why certain vaccines can induce reprogramming the immune response, including the innate immune responses, leading to nonspecific protection to reinfection and also a specific response in those patients with comorbidities (Arunachalam 2021, Forgacs 2021). The internal organization, or configuration, of a natural system at a given time t, will be modeled by a category K, called the configuration category at t: the objects of K represent the components of the system which exist at t, and the arrows of K (which we simply call links) represent the interactions between them around t that define the present organization of the system.

Having defined immunocompetence as a hyperstructure that emerges from the functional network that recognizes, neutralizes, and memorizes a given antigen, it is then necessary to ask about a possible mechanism that can reconfigure the state of the immune system of an individual and that allows him to pass from a deficient immune status to an immunocompetent one. our proposal is an immunocompetent adjunction, which we will explain below.

### The Immunocompetent adjunction

Let *Kvac* and *Kinf* be, respectively, the categories of vaccinated and infected status of an individual, configured respectively at *t0* and *t1*. An adjunction between *Kvac* and *Kinf* is a pair of functors

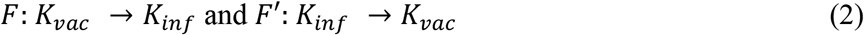

Together with a natural isomorphism α whose components for any objects V in *K*_*vac*_ and cP’ in *K*_*inf*_

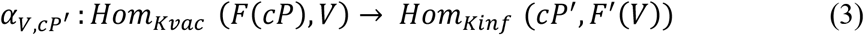

α, once the previously vaccinated person become infected, enables the activation of the immunocompetence hyperstructure of the configuration, built by vaccination at t_0_, to the configuration present at t_1_, allowing the generation or induction of a protective immune response in the individual. In brief and from a categorical point of view, we can propose that vaccines induce an adjoint functorial process, that optimally, reorganizes the configuration of the immune system of an individual leading to an effective immunocompetent state, which confers protection against the pathogen for which the vaccine was designed (**Figure 2**).

### Effective COVID-19 vaccines as transition drivers between immune configurations

Having defined immunocompetence as a colimit of an immune pattern/network, two implications arise. The first refers to the fact that an immune network induced by any antigen may or may not generate a state of immunocompetence against the antigen, which is a widely known fact in clinical immunology. Second, there is a possibility that different immunological patterns/immune networks, can generate an equivalent state or level of immunocompetence, which implies that the connectivity generated even by different sets of collective binding links, established between the effector cells of the immune system, plays a central role in establishing the immunocompetence status of an individual in response against a given antigen.

**Figure 2.**
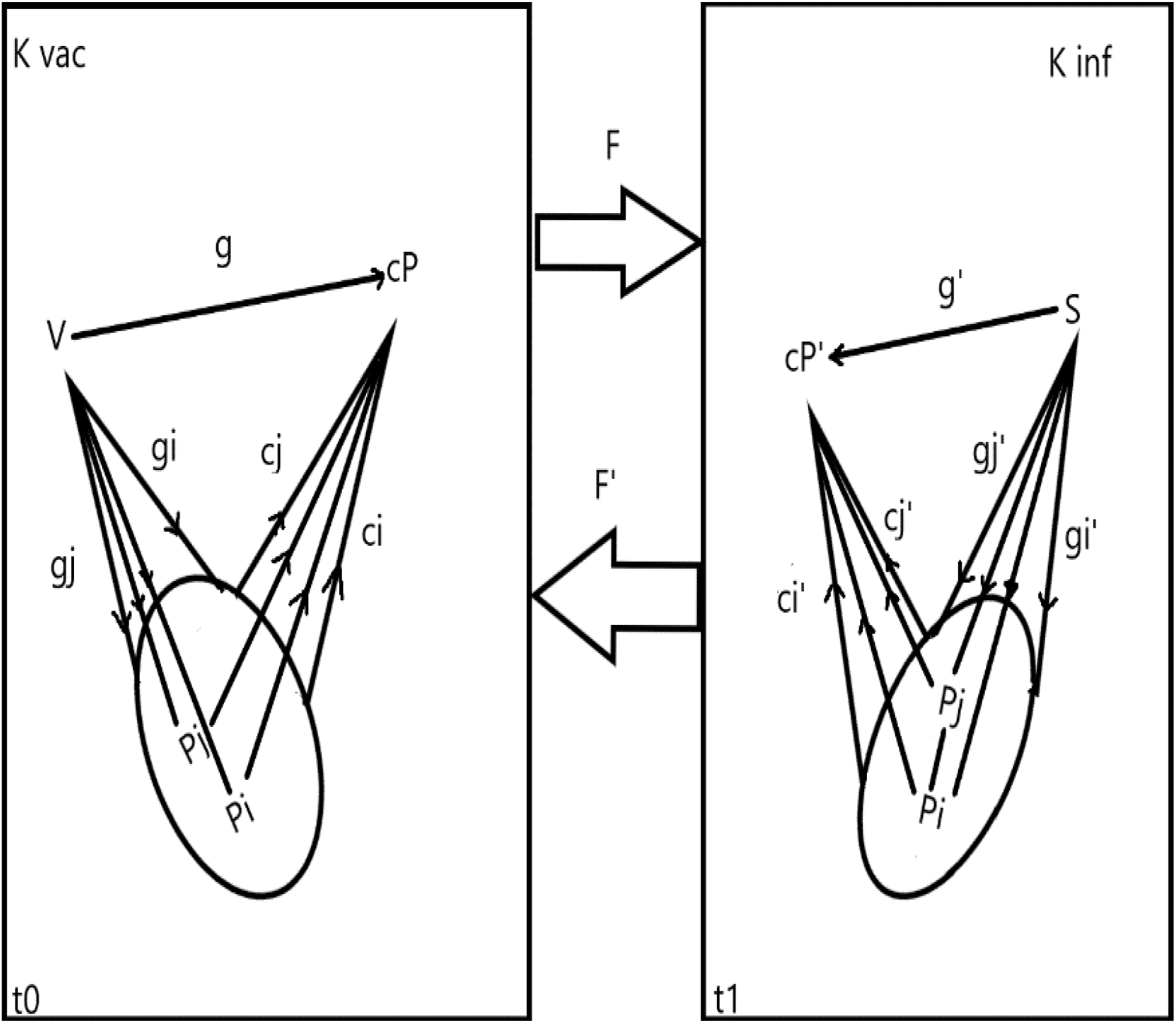
Reprogramming the immune response through an “adjoint like” process induced by effective vaccines. **Legend Figure 2**. The proposed mechanism for the action of an effective vaccine against a given pathogen can be described as a functorial reconfiguration, where at t_0_, the vaccine applied to an individual *Kvac*, generates, through its antigenic structure, a set of signals (*g*_*i*_) that induce the creation of a network of effector cells of the immune response (*P*_*i*_), which can include different aspects (*c*_*i*_), such as the innate immune response and also the adaptive immune response, configuring a higher-order immunoregulatory network (*cP*) or immunocompetence. In period t_1_, immune reconfiguration occurs when the same individual, vaccinated at t_0_ and potentially immunocompetent, is exposed to the target pathogen of the vaccine (*Kinf*), such as SARS-CoV-2, building an immune system state (*cP’*) which may or may not be protective or immunocompetent. The functorial mechanism (*F, F’*), previously expressed in equation (3), guarantees that the immunocompetence state generated at t_0_ by vaccination complements the state of the immune system at t_1_ by inducing immunoregulatory signals that end in an effective neutralization of the pathogen.

Recently, and constituting one of the most impressive technological achievements of the last decades and in response to the COVID-19 pandemic, a set of highly effective and safe vaccines against the SARS-CoV-2 coronavirus was developed and implemented in record time (Burgos 2021, Forni 2021). As its application expands to larger and larger human groups around the world, a series of very interesting facts begin to be documented.

First, many vaccinated individuals, even with a single dose and presenting comorbidities associated with severe cases of the disease, who become infected by the coronavirus, do not develop the disease or do so in a mild symptomatic way (Thompson 2021). Secondly, it has begun to detect the occurrence of activation of the innate immune response in vaccinated individuals, allowing them to react positively to the presence of other pathogens (Mckechnie 2020, Zhuofan 2021). Finally, it is observed that several of the vaccines against the original strain of SARS-CoV-2, regardless of their technological platform, confer protection to the current heterologous variants of concern (VOC) of coronavirus (Reynolds 2021).

These biological emerging facts are putting a growing interest in the notion of immunocompetence, given that immunologists have long known that, immunologically, more is not necessarily better. Moreover, how these immunological measures are obtained (usually in response to artificially administered antigens like vaccines) are not necessarily relevant. Rather, immune responses should be assessed by how effectively they protect an individual from infectious disease, given that this is the evolved function of the vertebrate immune system; that is, immunocompetence or immuno-incompetence should be measured and defined functionally rather than immunologically. Clinicians have been aware of the importance of functional measures for centuries; the first immunization strategies (for smallpox and human leishmaniasis) were based empirically on the ability of the immunization process to protect against subsequent disease, without any knowledge of either the nature of the infectious organism or the workings of the immune system (Bouwman 2020; Ganti 2021).

An interesting implication of the reconfiguration mechanism of the immune system (Figure 2), induced by an effective vaccine is that nothing prevents it from being carried out both in healthy patients and in patients with comorbidities and even infected patients (Arunachalam 2021, Schenten 2021). Normally this is not easy to demonstrate with commonly used vaccines such as polio, hepatitis B, or polysaccharides against meninge cocci, since a complete characterization of the individual’s immunocompetence status is not usually carried out before the application of the vaccine, even though the unquestionable successes of global vaccination campaigns against these pathogens seem to indicate that functorial reconfiguration may be quite common when effective vaccines such as the aforementioned are used.

On the other hand, and as a result of the development of excellent vaccines against SARS-CoV-2 and the application of transcriptomic analysis technologies, it is possible to carry out detailed studies of the effect of vaccination to control COVID-19, both in healthy patients and in a previously infected patient, finding that the vaccines, particularly RNA vaccines, induce an adequate state of immunocompetence in both types of patients, even when clear differences are observed in the response patterns, which supports our assertion that several patterns may have the same colimit, that is, concur in a state of immunocompetence that protects them against the same pathogen (Reynolds 2021).

### How to measure the immunocompetence level of an organism?

Having built a qualitative mathematical model, it is natural to ask, how can we verify the applicability of the model? since the structure underlying K is a graph and in particular, a random graph of finite order (*n* 〈∝) it is possible to estimate the probability of development of a state of immunocompetence in an organism under a given antigenic context, taking into account the following conditions:

1. Given that the basic property of a graph is to be connected, it is possible to assume that the functionality of the state of immunocompetence in an organism depends on the connectivity of the graph that represents the immune pattern *P* underlying *K*, that is, the limit *cP* exists if *P* is connected.
2. To estimate the probability that the graph that represents the immune pattern *P* conformed by a number *L* of links (*P*_*L*_), is connected, a fundamental theorem of Erdos and Renyi (1959) is used in which it is established that the probability that a small random graph is connected is related to the probability that it does not have vertices of degree 0.

#### Theorem 1.

let 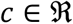 fixed and 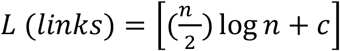 then *Prob*(*P*_*L*_ is connected)→ 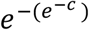. Where (*e*^*−c*^)= *λ*, corresponds with the mean percentage of isolated vertices in the immune pattern *P*_*L*_ following a Poisson distribution, and c represents a connectivity index where *c* = 0 implies that *P*_*L*_ is not connected.

To develop a practical case of measuring immunocompetence in a given organism, we will use one of the most interesting advances made in the study of the immune response mechanisms involved in the control of the SARS-CoV-2 infection in humans, the large-scale single-cell analyses of the immune characteristics of Covid-19 patients (Su 2020, Wilk 2020, Schmidt 2021). Single-cell RNA sequencing (scRNA-seq) is powerful at dissecting the immune responses under various conditions at the finest resolution and has been applied to COVID-19 studies by Ren and collaborators (2021) to a large cohort of 205 individuals, including hospitalized COVID-19 patients with moderate or severe disease, and patients in the convalescent stage, as well as healthy controls, observing critical changes in the peripheral blood discriminating mild/moderate from 147 severe COVID-19 patients in the disease progression or convalescence stages, form these data set, the authors count a total of 64 vertices for those signaling pathways associated with the adaptative immune response of severe COVID-19 patients, and 54 vertices for healthy patients (COVID-19 www.cellatlas.org).

Based on the information on the number of vertices of the immunological networks of patients with severe COVID-19 and healthy patients described by Ren (2021) and presented in Cellxgene (Cellxgene.cziscience.com/COVID-19.cancer-pku.cn/#marker-genes), using conditions 1 and 2 above and following Bollobas (2001), we carry out an estimate of the variation of the connectivity of the immune networks for different values of c and l, representing the result of antigenic stimulation in both types of patients, such as the effect of an effective vaccine against SARS-CoV-2. The results of the calculations of a putative dynamic of the connectivity of the immune networks of COVID-19 patients and healthy patients can be seen in Table 1.

**Table 1.** The probability of an individual of being immunocompetent against SARS-CoV-2 infection.**Legend Table 1.** The dynamics of immunocompetence induction are simulated, both in healthy individuals and with patients who have had COVID-19, assuming that both have received an effective vaccine against the coronavirus at t_0_ and that they subsequently infect or reinfect with the virus at t_1_. From our small random graph model of the immune pattern *P*, of order n = 54 vertices for healthy patients and n = 64 vertices for patients who have previously had COVID-19; it is noted that, as the connectivity index (*c*) increases, the probability that the graph is connected, which is interpreted as construction of immunocompetence, also increases rapidly in both types of patients, despite the initial difference in the order of the underlying graph. The calculations to obtain the data in this table are carried out using **Theorem 1** and the indications given in Bollobas (2001).

| Connectivity index | mean | Healthy patients |  | COVID-19 patients |  |
| --- | --- | --- | --- | --- | --- |
| C | $\lambda$ | Isolated vertices | probability | Isolated vertices | probability |
| 0 | 1 | 54 | 0,393 | 64 | 0,392 |
| 0,5 | 0,6 | 32 | 0,535 | 38 | 0,533 |
| 1 | 0,36 | 19 | 0,736 | 23 | 0,733 |
| 2 | 0,135 | 7 | 0,833 | 9 | 0,83 |
| 3 | 0,049 | 2,6 | 0,924 | 3 | 0,921 |
| 4 | 0,018 | 1 | 0,99 | 1 | 0,987 |
| 4,61 | 0,001 | 0,05 | 0,995 | 0,064 | 0,994 |

A first observation, arising from the data of the transcriptomic atlas, is related to the fact that the underlying immune networks of individuals who have suffered the disease and healthy individuals have a difference of an order of 15.6%, which would imply that in former patients a greater effort is required to achieve a state of immunocompetence against the coronavirus. On the other hand, the dynamics of increasing the connectivity index is exponential in both types of patients, as can be seen in Table 1, and despite the difference in the initial size of the immune networks considered, a state of connectivity is reached, exponentially, which would represent the emergence of immunocompetence in both types of patients as a result of vaccination.

## Conclusions

When modeling an organism, from the perspective of MES, as a category, whose immune system is represented as a P pattern made up of a network of effector cells and signals between them, generated as a result of their interaction with a given antigen, defines immunocompetence as emergent hyperstructure. A possible two-step mechanism of reconfiguration of the immune system induced by the application of an effective vaccine is proposed below. the first consists of the construction of a state of immunocompetence and the second, once in the presence of the pathogen, a process of optimization of the immune response networks of individuals is carried out, where any deficiency is supplied by immunoregulatory signals previously induced by effective vaccine application. In circumstantial or indirect support of the model, some recent results of vaccination with RNA vaccines against the coronavirus, that cause COVID-19 are considered, supporting the possible action of reconfiguration mechanisms of the immune system induced by the action of effective vaccines against SARS-CoV-2.

## Acknowledgements

The author fully acknowledges the support of CIINAS corp.

## Notes

### Competing Interest Statement

The authors have declared no competing interest.

